# Dual-axis Volta phase plate cryo-electron tomography of Ebola virus-like particles reveals actin-VP40 interactions

**DOI:** 10.1101/2021.04.21.440744

**Authors:** Sophie L. Winter, Petr Chlanda

## Abstract

Cryo-electron tomography (cryo-ET) is a pivotal imaging technique for studying the structure of pleomorphic enveloped viruses and their interactions with the host at native conditions. Owing to the limited tilting range of samples with a slab geometry, electron tomograms suffer from so-called missing wedge information in Fourier space. In dual-axis cryo-ET, two tomograms reconstructed from orthogonally oriented tilt series are combined into a tomogram with improved resolution as the missing wedge information is reduced to a pyramid. Volta phase plate (VPP) allows to perform in-focus cryo-ET with high contrast transfer at low-resolution frequencies and thus its application may improve the quality of dual-axis tomograms. Here, we compare dual-axis cryo-ET with and without VPP on Ebola virus-like particles to visualize and segment viral and host cell proteins within the membrane-enveloped filamentous particles. Dual-axis VPP cryo-ET reduces the missing wedge information and ray artifacts arising from the weighted back-projection during tomogram reconstruction, thereby minimizing ambiguity in the analysis of crowded environments and facilitating 3D segmentation. We show that dual-axis VPP tomograms provide a comprehensive description of macromolecular organizations such as nucleocapsid assembly states, the distribution of glycoproteins on the viral envelope and asymmetric arrangements of the VP40 layer in non-filamentous regions of virus-like particles. Our data reveal actin filaments within virus-like particles in close proximity to the viral VP40 scaffold, suggesting a direct interaction between VP40 and actin filaments. Dual-axis VPP cryo-ET provides more complete 3D information at high contrast and allows for better interpretation of macromolecule interactions and pleomorphic organizations.

**Highlights:**

- Volta phase plate dual-axis cryo-electron tomography provides high contrast tomography data with reduced back-projection ray artifacts and missing wedge information in Fourier space
- Volta phase plate dual-axis cryo-electron tomography facilitates interpretation of protein-membrane interactions
- Volta phase plate dual-axis cryo-electron tomography reduces ambiguity in manual 3D rendering and markedly improves 3D isosurface modeling
- Ebola virus-like particles contain actin filaments in close proximity to the VP40 layer

## Introduction

Cryo-electron tomography (cryo-ET) is an important imaging technique allowing *in situ* structural investigation of macromolecular organization in three dimensions (3D) and at close-to-native condition. A vitrified biological sample is tilted to acquire a set of projections at different angles which are aligned and back-projected to obtain three-dimensional information of the sample. Transmission electron microscopy is limited to thin samples that consequently must be shaped into sections or spread into a thin film placed on a meshed electron microscopy grid. As the sample is tilted the mean free-path of the electrons increases with increasing tilt resulting in a decreased signal-to-noise ratio at high tilts and limiting the angular range to ±60-70°[1]. Such incomplete angular sampling results in missing wedge information in Fourier space that in turn yields an anisotropic resolution of the reconstructed volume with elongation of features in Z-direction and a loss of information in the direction perpendicular to the tilt axis[2]. The angular resolution of the tomograms is defined by the Crowther criterion and depends on the number of projections and the thickness of the sample[3]. In order to partially restore the missing wedge information, dual-axis tomography can be performed[4]. In this method, two tilt series are acquired on the same area of interest that is rotated by 90° around the optical axis of the microscope. The two tilt series are independently aligned and reconstructed and the resulting tomograms are combined in the available software packages IMOD[5] or AuTom[6, 7] or the dual-axis tomogram is directly reconstructed from the orthogonal tilt series in SPIDER[8] or Protomo[9]. It has been shown that dual-axis tomography is beneficial to reduce reconstruction artifacts and to increase the amount of information in the 3D reconstructed volume. Dual-axis electron tomography is commonly applied in heavy metal stained resin-embedded thick-section samples that are besides shrinkage not sensitive to beam damage using either transmission[4] or scanning transmission electron microscopy[10, 11]. In cryo-electron microscopy, phase contrast dominates, and the total electron dose used per imaging area is limited due to the damage by electron radiation. To perform dual-axis cryo-ET the total dose per tomogram must be halved which in turn further decreases the signal-to-noise ratio and may result in inaccurate tomogram combination and contrast transfer function (CTF) determination of the individual projections. Nevertheless, several successful studies using dual-axis cryo-ET were conducted using microtubules[6, 7], polyribosomes[12], ATP synthase[13], and whole cells[14]. Owing to the development of direct electron detectors[15] and the Volta phase plate (VPP)[16], the signal-to-noise ratio in cryo-ET has substantially improved presumably facilitating dual-axis cryo-ET. However, dual-axis VPP cryo-ET has not been fully explored yet and only limited studies exist[17]. VPP is a hole-free phase plate composed of a thin amorphous carbon film (∼10 nm) placed in the back focal plane and heated to ∼200°C to prevent undesirable charging. In contrast to hole phase plates, VPP does not require accurate positioning, which makes its alignment user-friendly. As the zero-order beam passes through the carbon film of the VPP, an electrostatic negative potential is locally created on the surface of the carbon film, which in turn advances the phase of the unscattered wave by 90° without introducing a defocus[16]. This provides a phase modulation of the wave function, which is converted by the phase plate into an amplitude change in the image plane. As a result, the contrast between weakly scattering biomolecules and the surrounding vitrified environment is increased and the contrast transfer of low-resolution frequencies provided by VPPenables in-focus cryo-ET. VPP tomograms acquired with defocus target 0 µm have limited resolution to the first zero of the CTF. During tilting, the projection images exhibit a defocus gradient in the direction perpendicular to the tilt axis, leading to a gradient of achievable resolution dependent on the tilt and area of the sample[18]. This limitation can be circumvented by acquiring the VPP tilt series with a small defocus, which facilitates CTF determination and correction.

Cryo-ET has been pivotal in structural studies of pleomorphic viruses, such as Ebola virus (EBOV)[19-21]. EBOV belongs to the family of *Filoviridae* and forms several microns long filamentous particles that contain a negative-sense RNA genome[22] encapsidated by a helical nucleocapsid composed of the viral nucleoprotein N, VP24 and VP35 proteins[19, 20]. The nucleocapsids are incorporated into virus particles at the plasma membrane of infected cells, where the viral scaffold protein VP40 and the transmembrane glycoprotein GP are accumulated. VP40 is sufficient to drive virion budding and is responsible for the characteristic filamentous shape[23]. Recent cryo-ET studies revealed the organization of VP40 proteins in budded particles into an apparent helical scaffold lining the inner viral membrane leaflet[21]. Besides VP40 as a major driver of EBOV budding, actin has been shown to be indispensable, since the disruption of the actin cytoskeleton abrogates particle formation[24]. Actin co-localizes with VP40 at the periphery of the plasma membrane and appears to coordinate its movement[25]. However, how VP40 interacts with actin and whether actin is incorporated into budded particles remains unknown.

To investigate the structural components of EBOV and to evaluate the application of VPP in dual-axis cryo-ET we imaged Ebola virus-like particles (VLPs) VLPs[26, 27] that closely resemble the virus[19]. Here, we summarize the results and applicability of VPP dual-axis cryo-ET in direct comparison to single-axis and defocus phase contrast (DPC) cryo-ET. While the acquisition of VPP dual-axis cryo-electron tomograms is more laborious and time-consuming, it greatly facilitates the segmentation accuracy and analysis of components of virus particles from small datasets. This allowed us to directly visualize different nucleocapsid assembly states and to quantify the distribution of glycoproteins on the virion surface. We could further visualize and segment actin filaments in purified Ebola VLPs, which are found in close proximity to the VP40 layer.

## Results

To demonstrate the applicability of dual-axis cryo-ET to biological samples, we used Ebola VLPs and nucleocapsids because of their characteristic filamentous shape. The position of the filamentous particles in respect to the tilt axis determines the position of the missing wedge information within the filamentous particle. Filamentous particles positioned along the tilt axis are better resolved in X-Y and show lower elongation in the Z-direction than particles positioned perpendicularly to the tilt axis[4]. Ebola VLPs and nucleocapsids were produced by overexpression of the structural viral proteins in HEK 293T cells and isolated 48 hours post-transfection from cell media by sucrose-cushion ultracentrifugation. To acquire dual-axis VPP tilt series, we used a dose-symmetric schema with an angular range of ±60° and 3° increments with a total dose of 65 e^-^/Å^2^ per tilt series, which offers optimal electron dose distribution and minimal changes in microscope optics during tilt series acquisition[28] (Figure 1A). In comparison, unidirectional schemas starting at high angles significantly deteriorate the achievable resolution in the tomograms[28], while bidirectional schemas impact the VPP-induced phase shift as they require an additional realignment step during the tilt series that involves switching to low magnification. After a single title series was acquired, the stage was rotated using the Krios ‘rotate’ function (nominal rotation value of 97.3°). The area of interest was found back by using the navigator tool in SerialEM [29]. Tilt series were aligned, reconstructed and combined in IMOD (Figure 1B). Transverse tomographic cross-sections of a filamentous VLP oriented orthogonally to the tilt axis showed strong elongation of the VLP features in Z-direction, while the VLP appeared less elongated after stage rotation. The combined dual-axis tomogram exhibited more complete structural details of the particle membrane and the VP40 layer (Figure 1C).

**Fig. 1.**
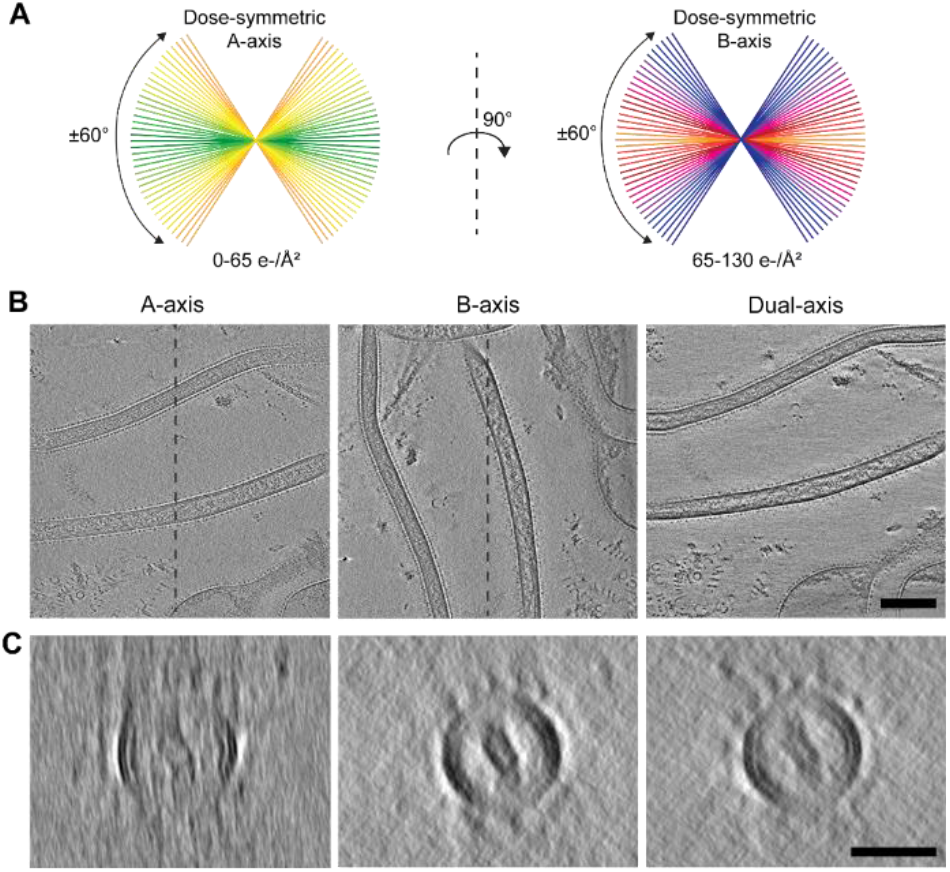
Dual-axis VPP cryo-ET using two orthogonal dose-symmetric tilt series acquisition schemas. A) Tilt series were acquired in-focus with VPP using dose-symmetric schemas with an angular range of ±60° and 3° increment around the sample axis with the total dose indicated below each schema. After the acquisition of the A-axis tilt series, the stage is rotated approximately 90° around the optical axis of the microscope and the B-axis dose symmetric tilt series is acquired. B) Slices of tomograms (8 nm thick) showing two filamentous VLPs orthogonally oriented to the tilt axis (A-axis), oriented parallel to the tilt axis after an approximate 90° microscope stage rotation (B-axis), and after the combination of A- and B-axis tomograms (Combined). C) Transverse tomographic slices (16 nm thick) of a filamentous VLP orthogonally oriented to the tilt axis (A-axis), oriented parallel to the tilt axis after an approximate 90° microscope stage rotation (B-axis), and after the combination of A and B-axis tomograms (Dual-axis). Scale bars: 100 nm (B); 50 nm (C).

### Comparison of tomogram combination in VPP and DPC dual-axis tomography

Critical for the combination of the single-axis tomograms is accurate tomogram registration. Therefore, we first evaluated whether the increased contrast of VPP tomograms facilitates cross-correlation based tomogram combination in IMOD. We used cross-correlation patches with different sizes to combine tomograms reconstructed from tilt series acquired with DPC (−4 µm defocus, which provides a relatively strong contrast transfer at low-resolution frequencies) or with VPP contrast at zero defocus. As expected, larger patches yielded higher cross-correlation values and better results in tomogram combination. Patches with dimensions of 400 × 400 × 200 pixels were required to obtain high cross-correlation values and to minimize the warping required to combine the tomograms (Figure 2E, F). Warping mean residual errors were similar in both DPC and VPP tomograms for different cross-correlation patch dimensions, indicating that increased contrast in low-resolution frequencies in VPP tomograms does not considerably improve the tomogram registration (Figure 2E, F). Since the dual-axis tomograms were acquired with a relatively high total electron dose of 130 e^-^/A^2^ at which the high-resolution details deteriorate, we next sought to evaluate dual-axis tomograms using a total dose of approximately 65 e^-^/A^2^. We could show that even at the total dose of 30-35 e^-^/A^2^ per single tilt series, the final tomogram combination was not impaired as apparent from the mean residual warping errors, which are similar to those obtained from tilt series acquired at higher dose (Figure 2F).

**Fig. 2.**
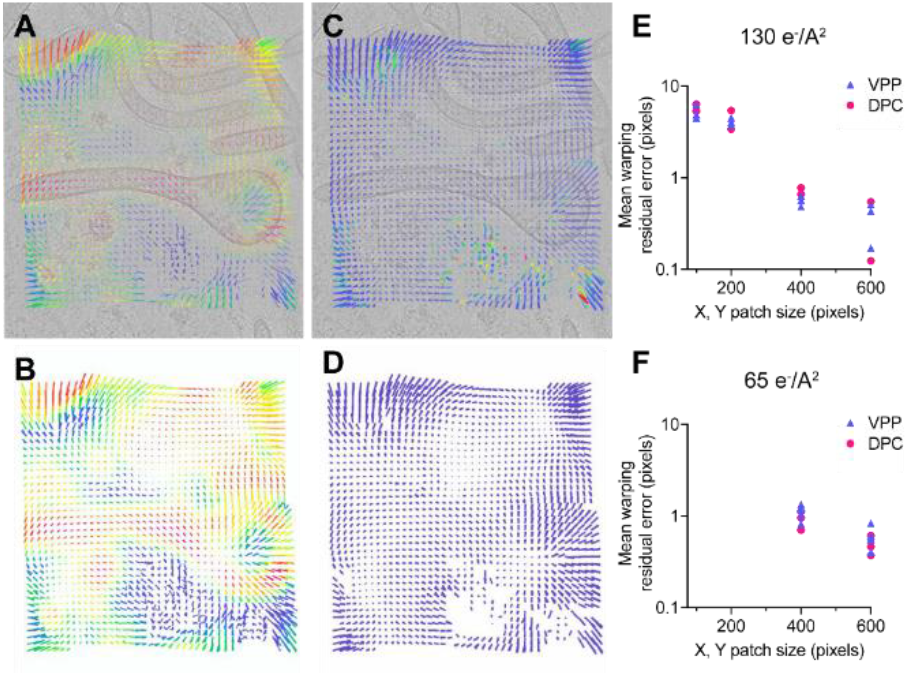
Tomogram combination using correlation patches in IMOD. Colloidal gold beads were used to perform an initial registration of the A-axis and B-axis tomograms. Subsequently, large overlapping patches (400 × 400 × 200) were used to refine the tomogram registration. A) Slice of a combined tomogram overlaid with vectors color-coded to represent the cross-correlation values (red - high cross-correlation values, blue - low cross-correlation values). B) Vectors representing cross-correlation values shown without the corresponding tomogram. C) Respective patch vector model representing the warping error overlaid with the slice of the tomogram. High errors are shown in red, low errors in blue. D) Patch vector model shown in C with high errors removed. The length of the vectors shows 10× the actual values. E) and F) Plots showing mean residual warping errors obtained in VPP and DPC dual-axis tomograms acquired at total electron doses of 130 (E) and 65 e^-^/A^2^ (F) using different patch sizes. The X-axes show the patch dimension in X and Y, with the Z dimension half of X.

### VPP dual-axis tomography improves visualization of protein-membrane interactions

We next evaluated the VP40 layer that directly interacts with the VLP membrane in single-axis and dual-axis tomograms reconstructed from projections using either DPC with a -4 µm defocus or VPP contrast. In all tomograms, the membrane bilayer lined with a VP40 layer at the lumenal side was visible inside the VLPs (Figure 3). The expected contrast boost was apparent in the VPP single-axis tomogram (Figure 3C, D) when compared to the tomogram acquired at -4 µm defocus (Figure 3A, B). A positive contrast (white halo) was prominent between the VP40 layer and membrane in the DPC tomogram, presumably caused by a fringing artifact arising from defocusing (Figure 3B). The in-focus VPP tomogram revealed a direct contact between the approximately 5 nm thick helically arranged VP40 layer and the membrane bilayer (Figure 3D) consistent with a recent subtomogram average[21]. The side-by-side comparison of DPC and VPP dual-axis tomograms showed that the VP40 layer and the membrane bilayer have higher contrast and reveal more structural details with reduced projection ray artifacts in the VPP dual-axis tomograms (Figure 3D). Dual-axis tomograms acquired with a total electron dose of 65 e^-^/A^2^ instead of 130 e^-^/A^2^ displayed markedly higher signal-to-noise ratio when VPP was used (Figure 3B). Hence, VPP dual-axis tomography provides sufficient contrast even at a highly reduced total electron dose.

**Fig. 3.**
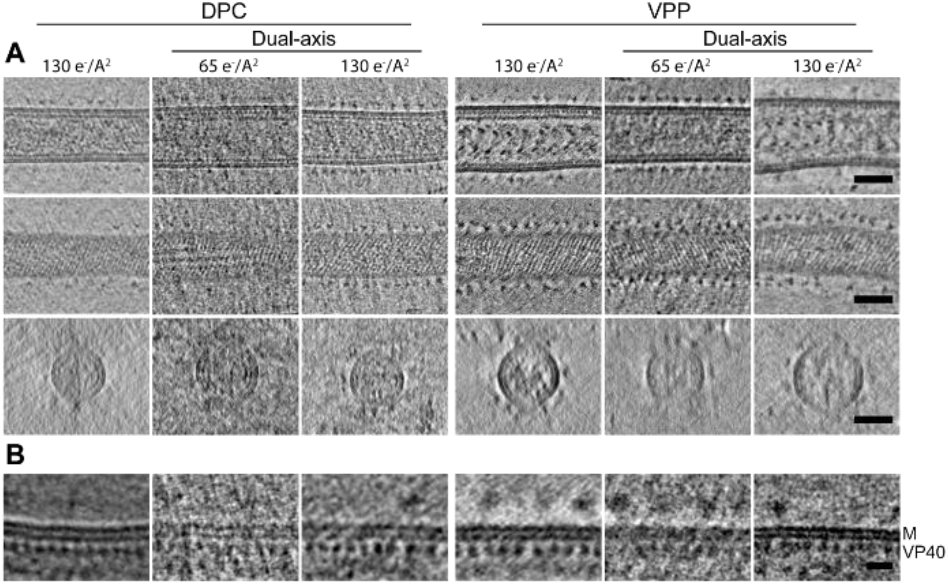
Comparison of dual-axis versus single-axis cryo-electron tomography using either DPC or VPP. Tomogram slices (8 nm thick) of Ebola VLPs reconstructed either from single or dual-axis tilt series collected with VPP in-focus or without VPP at defocus -4 µm. Tilt series were acquired with a total dose of either 65 e^-^/A^2^ or 30-35 e^-^/A^2^. A) Slices of tomograms showing the longitudinal central slice (top), longitudinal near-to-surface slice showing the striation of the VP40 layer (center), and transverse slice (bottom) through a VLP. B) Slices through tomograms showing the membrane (M) and VP40 layer. Scale bars 200 nm (A); 50 nm (B); 10 nm (C).

### VPP dual-axis tomography reduces Z-elongation and provides high contrast imaging of complex assemblies

We next evaluated the reduction of the missing wedge information in the dual-axis VPP data. Closer examination of 10 nm protein A coated colloidal gold clusters that are used as fiducial markers during tilt series alignment revealed a reduction of the missing wedge information in dual-axis tomograms. The spherical shape of the gold bead was better represented in dual-axis tomograms (mean Z-elongation 1.32), whereas the gold bead appeared more elongated in the Z-direction (mean Z-elongation 1.75) in the single-axis tomogram, which is in agreement with previous studies[4]. Moreover, the back-projection ray artifacts orthogonal to the tilt axis apparent in single-axis tomograms in X-Y were markedly reduced in dual-axis tomograms (Figure 4).

**Fig. 4.**
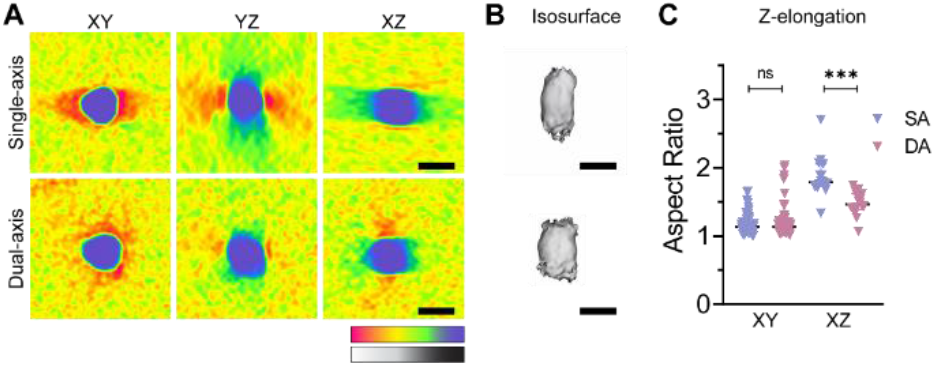
Single-axis versus dual-axis VPP cryo-ET of colloidal gold beads. A) Orthogonal tomogram slices (X-Y, Y-Z, X-Z) through the center of one gold bead. False-color map used instead of greyscale for visualization (color key displayed below the images. B) Isosurface of gold bead shown in (A). C) A scatter plot showing means and 95% confidence intervals of aspect ratios of measured gold bead diameters in X-Y (n = 57) and X-Z (n = 16) in single- and dual-axis tomograms. Unpaired two-sided *t*-test showed a significant reduction in the Z-elongation of particles measured from DA compared to SA tomograms (p<0.0001). Scale bars: 10 nm.

To further assess the reduction of the missing wedge information in the dual-axis VPP tomograms on biological structures, we next investigated EBOV nucleocapsid helical assemblies. Ebola VLPs predominantly contained loosely coiled nucleocapsids, which was particularly obvious in VPP tomograms (Figure 3). Loosely coiled nucleocapsid assemblies were also found outside of the VLPs, likely released from the overexpressing cells after their lysis or from broken VLPs (Figure 5A). To examine the reduction of missing wedge information in dual-axis VPP tomograms, we used isosurface 3D modeling of the same area containing nucleocapsid assemblies in single-axis or dual-axis tomograms. The reduction of missing wedge information was particularly apparent in the isosurface representation of the loosely coiled nucleocapsid forming a solenoid. In the dual-axis tomogram, the nucleocapsid solenoid structure appeared more complete, whereas in the single-axis tomogram the solenoid structure was elongated in Z-direction and contained gaps in density (Figure 5B). In addition to the loosely coiled nucleocapsid that had an approximate diameter of 35 nm, two other forms of nucleocapsid assemblies representing a condensed nucleocapsid and condensed nucleocapsid decorated with additional protein densities were observed outside of the VLP (Figure 5C). We thus demonstrate that VPP dual-axis tomography provides near complete 3D information with high signal-to-noise ratio.

**Fig. 5.**
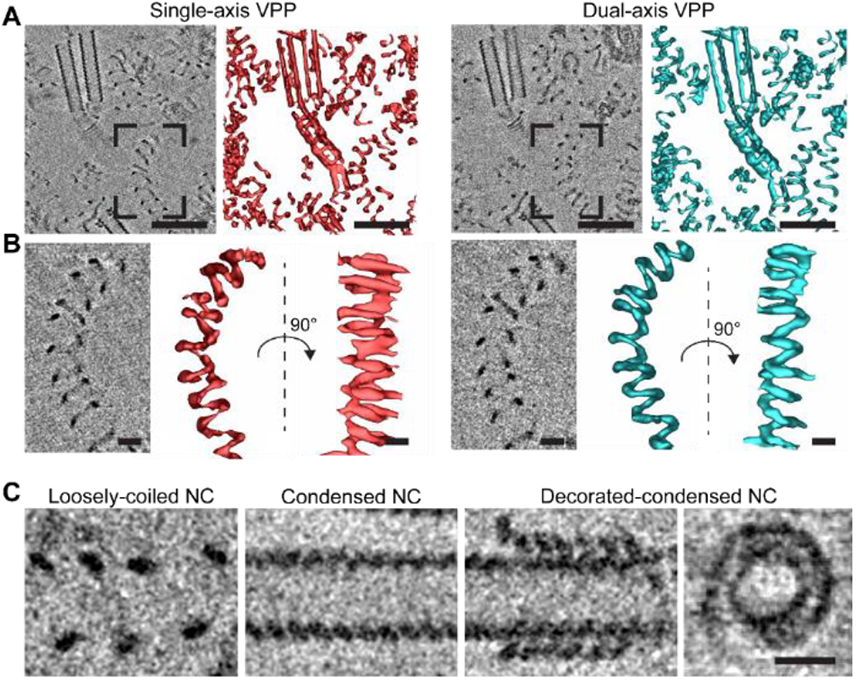
VPP single- and dual-axis tomography of EBOV nucleocapsid assemblies. A) Slices of tomograms (8 nm thick) showing Ebola nucleocapsids either reconstructed from a single-axis dose-symmetric tilt series with VPP (left) or from a dual-axis tomogram (right). B) Slices of tomograms (8 nm thick) capturing a loosely coiled nucleocapsid highlighted by a black frame in A. Corresponding isosurfaces of the densities found in the single and dual-axis tomograms are shown in red (single-axis tomogram) and in cyan (dual-axis tomogram). C) Slices of a dual-axis tomogram (8 nm thick) showing slices through a loosely coiled nucleocapsid, a condensed nucleocapsid, and a condensed nucleocapsid decorated with additional densities presumably corresponding to VP24 and VP35. Scale bars: 100 nm (A); 20 nm (B, C).

### VPP dual-axis tomography facilitates 3D isosurface modeling

We next evaluated the isosurface 3D modeling in DPC and VPP dual-axis tomograms using GP trimers protruding from the VLP surface. While in dual-axis DPC data the typical trimer organization was less obvious (Figure 6A-C), dual-axis VPP tomography provided a clear visualization of the EBOV GP trimers protruding from the VLP envelope, which closely resembled the subtomogram average of the GP spike[30]. Isosurface modeling of the GP trimers on the surface of the VLPs allowed straightforward 3D visualization and segmentation of the trimers for further analysis. Quantification of the distances between individual GPs revealed and average GP-GP distance of 15.3 nm (standard deviation (SD) 3 nm; n = 160).

**Fig. 6.**
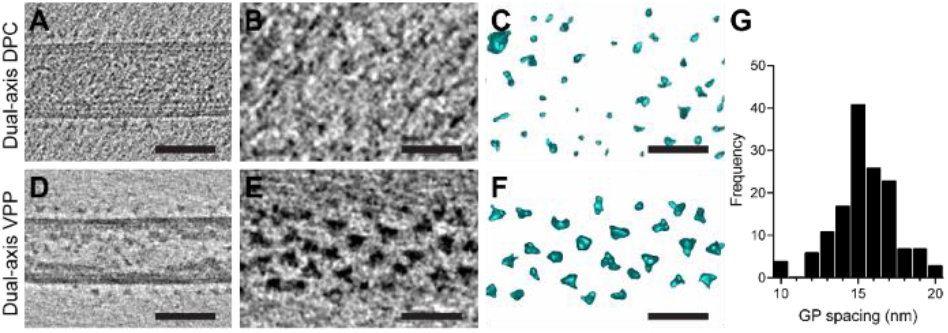
3D isosurface modeling of Ebola glycoproteins. Slices of tomograms (8 nm thick) acquired by dual-axis DPC (A-C) or dual-axis VPP cryo-ET (D-F). A) and D) show a longitudinal central slice of a VLP. B) and E) show top slices revealing the GP trimers, which are shown as isosurface rendering in (C) and (F). G) Histogram of GP-GP distances in nm calculated from coordinates of GP proteins segmented from the VLP in dual-axis VPP data shown in D. The average GP-GP spacing was 15.3 nm (SD 3 nm; n = 160). Scale bars: 50 nm (A, D); 20 nm (B, C, E, F).

### VPP dual-axis tomography facilitates manual volume segmentation of the VP40 layer actin filaments inside VLPs

We frequently observed filamentous structures in the proximity of the VP40 layer in VLPs (Figure 7A, B, D, E) and occasionally found structurally similar filaments outside the VLPs, probably co-purified from the supernatant of VLP-producing cells (Figure 7C, F). In both cases, the filaments showed a diameter of approximately 7 nm (n= 20 isolated filaments, n= 13 filaments in VLPs) and were composed of helically organized subunits consistent with F-actin filaments (Figure 7C). The filaments were exclusively observed directly underneath the VP40 layer (Figure 7B, H, I) following the tubular virus particle shape, and appeared to be connected to VP40. VPP dual-axis tomography facilitated manual segmentation of the actin and VLP components in Amira and enabled their three-dimensional visualization (Figure 7H, I). We could thereby visualize the global VP40 organization into a helical scaffold directly underneath the membrane. Additionally, we observed the VP40 scaffold at areas, where the filamentous VLP morphology is altered including bifurcations and where the particle is bending. To visualize the VP40 arrangement in these regions we rendered areas of VLPs with non-filamentous morphology captured in dual-axis VPP tomograms (Figure 7J-L). VLPs with globular morphology showed patches with linearly arranged VP40 oligomers without a clear helical symmetry that were interspaced with gaps lacking VP40 (asterisks, Figure 7K) and discontinuous linear VP40 oligomers (arrowheads, Figure 7L). This suggests that globular shape is determined by the high flexibility of the VP40 lattice that accommodates different shapes by gaps and patches of linearly arranged VP40 dimers.

**Fig. 7.**
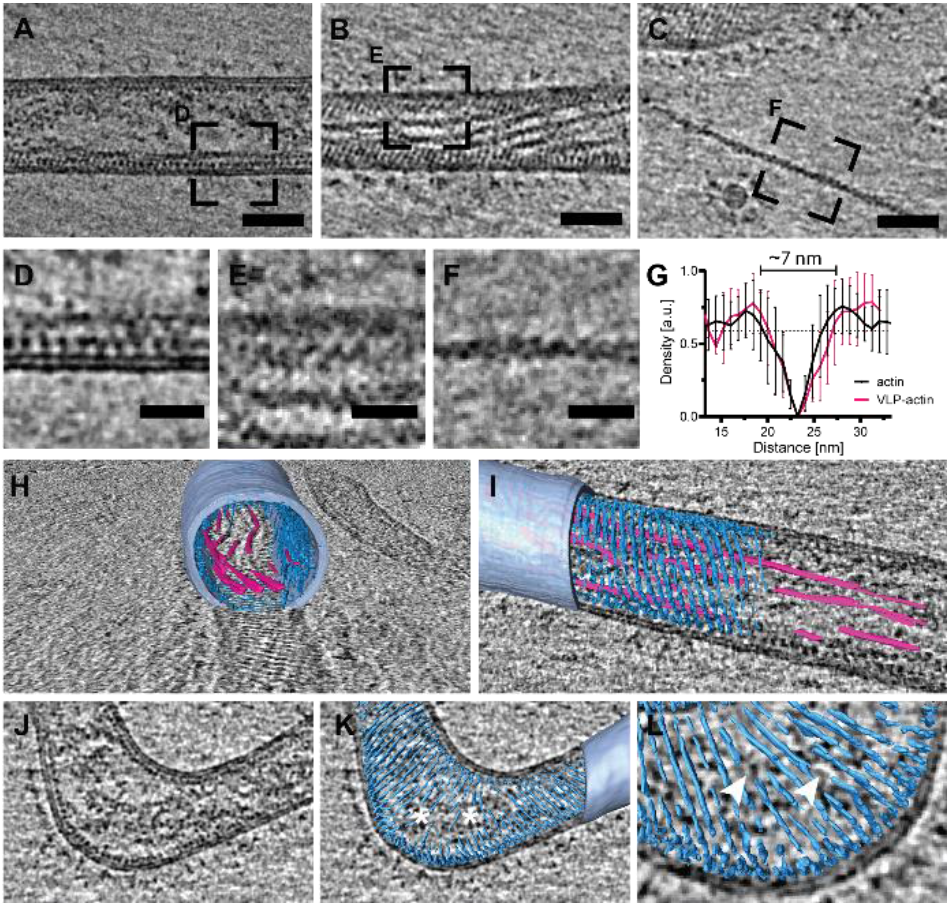
Ebola VLPs contain actin filaments in close proximity to the VP40 layer. A) Longitudinal central slice of a dual-axis VPP tomogram (8 nm thick). B) Longitudinal near-to-surface slice showing the striation of the VP40 layer and actin filaments in close proximity. D, E) Respective details of the areas marked in A and B. Slices shown in B-F are 4 nm thick. C) Slices through a tomogram showing an actin filament outside of a VLP and its detail in F. G) Normalized average line density profiles showing the diameter of the actin filament inside and outside of the VLP. The standard deviation is indicated for each measurement (n= 20 for actin outside a VLP, n= 13 for actin within a VLP). The average background grey value is indicated by dotted lines (magenta and black, respectively). H and I) Rendering of the VLP containing actin filaments (VP40 layer - cyan, actin - magenta, membrane - light blue). J-L) Rendering of the VP40 scaffold at a bent VLP area. Gaps in the scaffold are indicated by asterisks, discontinuous linear VP40 oligomers are marked with arrowheads. Scale bars 50 nm (A-C); 20 nm (D-F).

## Discussion

VPP cryo-ET is instrumental in the interpretation of tomograms thanks to the increased contrast at low-resolution frequencies in close-to-focus projections. Cryo-ET with VPP has been applied to study pleomorphic viruses and their interactions with membranes[31, 32] as well as to study the axonemes in trypanosome flagella[33]. In addition, VPP data is suitable for high-resolution studies. Single-particle analysis using either in-focus or defocused VPP projections of a 20S proteasome led to a resolution below 3.2 Å[34] and 2.4 Å[35], respectively, and subtomogram averaging of 80S ribosomes led to a resolution of 9.6 Å[18]. However, subtomogram averaging of the gag protein in immature HIV virions showed that higher resolution can be reached without VPP in a benchmarking study where several acquisition conditions were tested side-by-side [36]. The current limitations of the VPP are the changing phase shift during tilt series acquisition and the signal attenuation as about 10% of the electrons are absorbed due to the presence of the carbon film. A recent study shows that signal is reduced to ∼40% of that observed without a VPP at ∼2 Å resolution[37].

Nonetheless, VPP offers advantages for dual-axis cryo-ET, since it allows in-focus imaging and a transfer of contrast at low-resolution frequencies resulting in strong contrast even if the total dose per tilt-series is low. As expected and shown for DPC dual-axis tomography[4], dual-axis VPP tomograms containing colloidal gold beads and EBOV nucleocapsids showed less Z-elongation and more complete three-dimensional information when compared to single-axis tomograms. Qualitative comparison of dual-axis tomograms acquired with or without VPP showed that protein-membrane interactions are better represented in VPP dual-axis tomograms. The ray back-projection artifacts and the fringing were minimized in close-to-focus VPP dual-axis tomography, which allowed visualization of the VP40 layer in direct contact with the membrane. However, the increased contrast in VPP tomograms does not improve the tomogram combination, presumably because the low-resolution frequencies do not significantly contribute to the cross-correlation coefficient. In both VPP and DPC cryo-ET dual-axis tomography, a small amount of warping was required to combine the tomograms. Notably, similar results were obtained in tomograms acquired at lower electron dose indicating that beam-induced damage[38-40] is not the dominant factor affecting the tomogram combination in our setup at a higher dose.

To demonstrate the potential of dual-axis VPP cryo-ET when studying heterogeneous, biological samples, we focused on different components of purified Ebola VLPs. Dual-axis VPP cryo-ET greatly facilitated 3D isosurface modeling of glycoproteins on VLPs and allowed straightforward quantification of their distribution on the particle surface. Furthermore, it allowed 3D isosurface modeling of the helical nucleocapsid and its different condensation states including the loosely coiled and condensed decorated nucleocapsid with additional proteins consistent with a previous study[19]. Notably, the boomerang-shaped density of the decorated nucleocapsid was directly visualized in the dual-axis tomograms and is consistent with NP-VP25-VP35 interacting proteins[20]. Interestingly, unlike in previous studies, we also observed a condensed nucleocapsid composed only of NP proteins with a diameter of approximately 30 nm. Importantly, dual-axis VPP tomography allowed us to manually segment the VP40 scaffold and actin filaments in VLPs due to more complete 3D information along with the improved signal-to-noise ratio. The close proximity to the VP40 layer and lack of additional protein densities are suggestive of a direct interaction of actin with VP40. To date, the requirement of actin for budding of Filoviruses has been demonstrated by the efficient inhibition of particle formation in the presence of inhibitors of actin polymerization and actin motor proteins[41]. Co-localization of VP40 with actin has been shown by fluorescence light microscopy[24, 25, 42] and traces of actin could be observed by Western blotting in purified virus particles[24, 42]. Whether actin energetically contributes to membrane bending and protrusion of the long, filamentous particles from the plasma membrane in addition to its requirement for the transport of viral proteins to budding sites has been subject of debate. The direct visualization of actin filaments in Ebola VLPs imaged in this study provides further proof that VLPs incorporate actin and that VP40 might directly interact with actin filaments, without additional adaptor proteins. The number of filaments per particle and their persistence length that can be measured more reliably using dual-axis VPP cryo-ET as demonstrated here could further be used to calculate the contribution of actin to membrane bending.

A major disadvantage of using VPP dual-axis cryo-ET is the doubled microscopy acquisition time. In our system, a single-axis tilt series takes approximately 34 minutes. The new generation of single-axis holders available on the latest Thermo Fisher Scientific systems have improved mechanical performance and support faster tilt-series acquisition[43, 44] but do not allow for grid rotation. Therefore, a system with a free grid rotation inside the autoloader or in the Autogrid gripper in combination with a single-tilt holder would be desirable for faster dual-axis tomography. The variable conditioning time required to obtain a 90° phase shift and its instabilities could be partially mitigated by assessing the phase shift for each conditioning spot before starting a tilt series and during tilt series acquisition. Maintenance of a desirable phase shift during tilt series acquisition may be more challenging with direct electron detectors with slower readout. Recent advances in laser phase plate technology[45, 46] that provide a tunable and constant phase shift for in-focus cryo-EM hold great promise. Finally, dual-axis VPP tomography would be beneficial to interpret crowded cellular environments in cryo-lamellae. An optimized milling strategy permitting dual-axis tilting on cryo-lamellae needs to be established, and the use of VPP on lamellae requires some additional considerations such as platinum sputtering on lamella prior to data acquisition to avoid charging. Furthermore, the thick organometallic platinum protective layer deposited during the preparation of cryo-lamellae prevents using most of the grid areas as a site for VPP activation. Consequently, some areas of lamella must be sacrificed for VPP activation, which may cause lamella to destabilization. The applicability and benefit of VPP cryo-ET on lamellae has been demonstrated in a study of the nuclear environment of HeLa cells[47].

In summary, dual-axis VPP shows several improvements in tomogram quality and is a valuable tool for cryo-ET applications. It is particularly useful in cases where subtomogram averaging is not straightforward because of the small size and low abundance of the protein complexes of interest, or *in cellulo* where heterogeneous interactions between macromolecules are studied[48]. In such instances, VPP cryo-ET provides contextual information that can be complemented by other structural biology approaches.

## Material and Methods

### Cells and plasmids

HEK 293T cells were maintained in DMEM medium supplemented with 10% FBS for the production of Ebola VLPs. Cells were transfected with equal amounts of plasmids encoding the EBOV proteins GP, VP40, N, VP24 and VP35 and supernatants were harvested 48 hours post transfection. After two successive clarification steps at 1,500 ×g for 10 min and 3,500 ×g for 15 min, the supernatants containing the VLPs were centrifuged for 2.5 hours at 11,400 rpm (SW32 rotor, Optima L-90K ultracentrifuge (Beckmann)) through a 30% sucrose cushion (HNE buffer, 10 mM HEPES, 100 mM NaCl, 1 mM EDTA, pH 7.4). Pellets were resuspended in HNE buffer, combined and centrifuged again at 19,000 rpm (TLA 120.2 rotor, Optima TLX ultracentrifuge (Beckmann)). The final pellet was resuspended in HNE buffer and VLPs were immediately plunge frozen.

### Plunge freezing

VLPs were plunge frozen in liquid ethane kept at - 183°C using GP2 (Leica). A suspension containing a mixture of 10 nm protein A coated colloidal gold (Aurion) and Ebola VLPs was applied with a volume of 3 µl on a glow discharged EM grid (Quantifoil, 200 Mesh, R2/1). The grid was blotted using No.1 Whatman paper for 3 sec from the back-side in a chamber with controlled humidity set to 70% and the temperature set to 24°C.

### Cryo-electron tomography

Krios TEM (FEI, now ThermoFisher Scientific, USA) was operated at 300 kV in nanoprobe mode at magnification 33.000× (pixel size 2.671 Å) using 50 µm C2 aperture. Projections were acquired using the Quantum energy filter (Gatan, USA) with a slit 20 eV and K3 direct electron detector operating in counting mode. Tilt series were acquired using the dose-symmetric tilt schema with an angular range ±60°, an increment of 3°, and a projection dose of ∼1.5-3.2 e^-^/Å^2^ using SerialEM 3.8.0 beta [49]. Each record projection was acquired as a movie composed of 10-15 frames which were aligned and summed on-fly using the SerialEM SEMCCD plugin. Beam-tilt angle for focus determination was set to 10 mrad and no defocus offset was used. Beam shift pivot points were aligned for both microprobe and nanoprobe modes. Objective astigmatism correction and coma-free alignment were performed in Sherpa 1.12.3 (ThermoFisher Scientific, USA). An objective aperture with a diameter of 100 µm was used for the defocus phase-contrast cryo-ET. The VPP on-plane position at the back focal plane and the C2 condenser astigmatism was corrected using a Ronchigram with 150 µm diameter C2 aperture. VPP conditioning was done at a magnification 33.000×, beam diameter ∼2.5 µm and screen current ∼0.3 nA, and the phase shift of 0.5 π was verified using the Sherpa tool 1.12.3 before starting each tilt series acquisition.

### Tomogram reconstruction and combination

Tilt series were aligned using 30-40 fiducial markers. The CTF was corrected by phase flipping in tilt series collected without VPP, no CTF correction was done in VPP tilt series. Electron dose filtering was done using the Mtffilter program with default parameters (a = 0.245, b = −1.665, and c = 2.81) previously determined [50] and tomograms were reconstructed by weighted back-projection using the SIRT-like filter (10 iterations) in Etomo (IMOD version 4.10.49) [5]. Two tomograms were combined in IMOD using fiducial markers on one side, fiducial markers with high residual errors were excluded to keep the mean residual error below 8 pixels. The final rotation between the axis of the two tomograms was approximately 87°. Local cross-correlation values of overlapping patches (400×400×200 pixels) were calculated using kernel filtering with sigma 3, and linear interpolation for both initial registration and the final match. Large residual patch vectors corresponding to areas with low cross-correlation values were removed from the patch vector model to minimize the warping residuals. Tomograms were binned 3× for figure production. Tomogram isosurfaces were performed in IMOD (Figures: 4, 5, 6) and renderings (Figure 7) in Amira.

### Determination of GP spacing on the VLP surface

GP trimers were manually segmented on the VLP surface from dual-axis tomograms acquired with the VPP. Respective 3D coordinates were extracted and for each GP trimer, the neighboring GP in closest proximity was identified. The Euclidean nearest-neighbor distance was calculated using the pandas (https://pandas.pydata.org/pandas-docs/stable/whatsnew/v1.0.4.html) and numpy software libraries [51].

### Measuring Z-elongation of gold particles

To determine the Z-elongation of gold particles in single- and dual-axis tomograms acquired using the VPP, X-Y and Y-Z projections capturing gold particles in-focus were analyzed in FIJI/ImageJ. Using the default threshold, the gold beads were automatically segmented, and the aspect ratio for each individual fold bead was measured.

### Measurement of filament diameter inside and outside Ebola VLPs

Line density profiles (10 pixels in width) of ∼ 25 nm in length were determined perpendicular to the filaments using the plot profile tool in FIJI/ImageJ. For filaments outside and inside a VLP several measurements were performed (n= 20 and n= 13, respectively) and normalized against the maxima of each individual measurement. Normalized gray value profiles were aligned to their maxima, plotted against the distance from the start of the line profile and averaged. The filament diameter was defined as the distance between the intersections of the averaged, normalized line profiles with the background baseline corresponding to the average gray values of the sample environment.

## Acknowledgments

We thank Dr. David Mastronarde for advice on tomogram combination in IMOD and for critical reading of the manuscript. We thank Dr. Ronald N. Harty for providing us with the EBOV plasmids. We would like to acknowledge access to the infrastructure and support provided by the Cryo-EM Network at the Heidelberg University (HD-cryoNet). Funding: This work was supported by a research grant from the Chica and Heinz Schaller Foundation (Schaller Research Group Leader Programme) to PC and SLW.

## CRediT authorship contribution statement

Conceptualization: PC; methodology, SLW, PC; validation, SLW, PC; formal analysis, SLW, PC; investigation, SLW, PC; writing – original draft preparation, SLW, PC; writing – review and editing, SLW, PC; visualization (figures), SLW, PC; supervision, PC. funding acquisition, PC. All authors have read and agreed to the published version of the manuscript.

## Notes

### Competing Interest Statement

The authors have declared no competing interest.

